# Robust single membrane protein tweezers

**DOI:** 10.1101/2023.01.15.524119

**Authors:** Seoyoon Kim, Daehyo Lee, W.C. Bhashini Wijesinghe, Duyoung Min

## Abstract

Single-molecule tweezers, such as magnetic tweezers, are powerful mechanical manipulation tools that can probe nm-scale structural changes in a single membrane protein under force. However, the weak molecular tethers used in the tweezers limit a long time, repetitive mechanical manipulation because of their force-induced bond breakage. Here, using the metal-free click chemistry of dibenzocyclooctyne (DBCO) cycloaddition and the rapid, strong binding of traptavidin to dual biotins (2xbiotin), we developed robust single-molecule tweezers that can perform thousands of force applications on a single membrane protein. By applying up to 50 pN for each cycle, which is sufficiently high for most biological processes, we were able to observe repetitive forced unfolding for a designer membrane protein up to approximately 1000 times on average. Monte Carlo simulations showed that the average error of the unfolding kinetic values rapidly decays to 1.8% at 200-time pulling, indicating that our method can quickly produce reliable statistics. The approach established here is also applicable to highly polar DNA molecules, permitting the nanomechanical manipulation of diverse biomolecular systems.

## Introduction

Single-molecule tweezers are an emerging tool for membrane protein studies, which has yielded important insights into membrane protein biogenesis, interaction with drugs, and chaperone effects on their folding^1-3^. To apply mechanical force to a target protein in magnetic or optical tweezers^4-7^, we tether two protein positions, *e*.*g*., the N- and C-termini, to solid supports of a micron-sized bead and the sample chamber surface (or another bead), respectively. Rapid noncovalent tethering has been widely used, such as the binding between digoxigenin (dig) and antidigoxigenin (antidig), 6xHis tag and Ni-NTA, and biotin and streptavidin^8-22^. However, the noncovalent tethers are dissociated in tens-of-pN force ranges^23,24^, restricting the utilization of the tweezer methods.

In covalent tethering approaches developed for single-molecule tweezers (*e*.*g*., HaloTag-chloroalkane, thiol-maleimide, *etc*), at least one end of a protein or its tandem repeat is attached to a solid support without using long molecular spacers, such as DNA handles^25-32^. In this case, the close proximity to the large-area surface and/or between the consecutively-connected proteins of the tandem repeat can hinder membrane protein studies since the lipid bilayer mimetics such as bicelles or vesicles of ∼100 nm or larger in diameter will not be properly accommodated in the narrow space^19,33^. On the other hand, single-molecule tweezers using hundreds-of-nm long molecular handles flanking a protein of interest still rely on the conventional noncovalent tethers of the dig-antidig and biotin-streptavidin^10-15,19^. The dig-antidig interaction is as very weak as *τ*_off_ = ∼1 sec at 25 pN (∼0.1 sec at 50 pN)^24^, limiting the stable performance of experiments, data throughputs (tens of pulling on one molecule at most), and reliable statistical analyses.

Here, we describe a robust magnetic tweezer approach that can perform thousands of repetitive mechanical manipulations on a single membrane protein. To establish the tweezer approach, we employed three orthogonal tethering strategies: the metal-free DBCO-azide click reaction, strong binding between traptavidin (a streptavidin variant with 17-fold slower biotin dissociation and 8-fold stronger binding) and 2xbiotin, and spontaneous isopeptide bond formation between SpyTag and SpyCatcher. Our method was also successfully tested on a DNA hairpin with the opposite polarity, ensuring its wide applicability to diverse biomolecular interactions. The single-molecule tethering schemes developed here can be adopted in optical tweezers, the other widely employed tweezers, and all other versions of magnetic tweezers.

## Results

Our single-molecule tweezer approach is illustrated in Figure 1. The step-by-step procedure of the single-molecule tethering is shown in Figure 2: the surface modification of magnetic beads (Figure 2A), preparation of DNA handles (Figure 2B), passivation of sample chamber surface (Figure 2C), and final assembly of single-molecule pulling system (Figure 2D). To test the method, we adopted two biomolecules that are extremely different in polarity: a designer membrane protein with four transmembrane (TM) helices (scTMHC2)^34^ that is highly nonpolar, and a DNA hairpin with 17-bp stem and 6T loop (17S6L) that is highly polar (Figure 1). scTMHC2 is conjugated to 1024-bp long DNA handles at its N- and C-terminal ends^35^ (Figure 1–figure supplement 1). The DNA handles with a total 700 nm contour length secure enough space to accommodate an approximately 100 nm bicelle in diameter, which is a lipid bilayer disc wrapped in deteregents^19,33^. Reconstituting the lipid bilayer environment is essential to maintain the native fold and stability of membrane proteins^36^.

**Figure 1.**
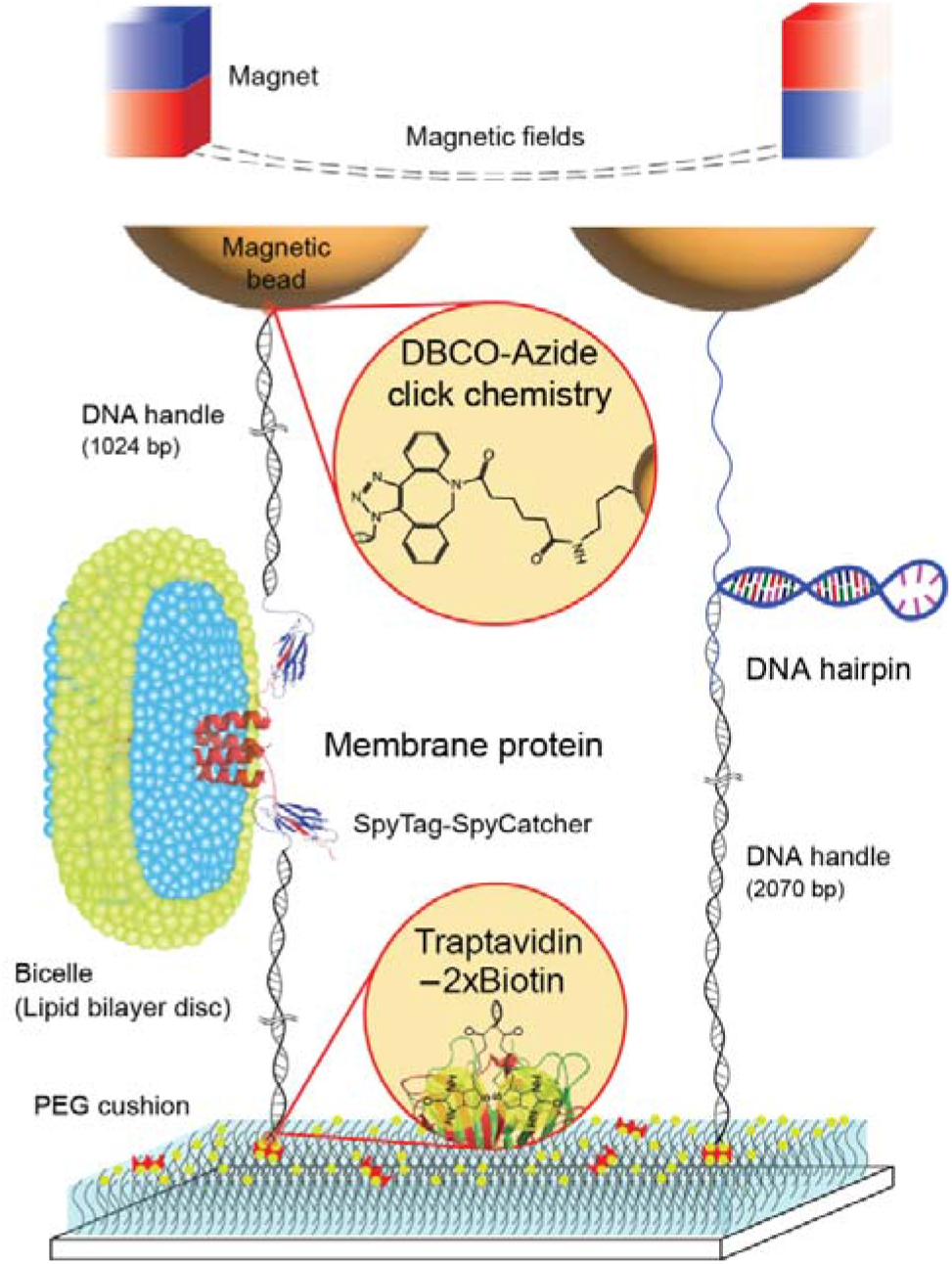
Schematic diagram of our robust single-molecule tweezers. Two biomolecules of a membrane protein, scTMHC2, and a DNA hairpin, 17S6L, were adopted to test the tweezer approach. DNA handles are attached to the membrane protein via spontaneous isopeptide bond formation between SpyTag and SpyCatcher. A DNA handle is attached to the DNA hairpin via annealing and ligation. One end of the molecular constructs is tethered to magnetic beads via DBCO-azide click conjugation. The other end of the constructs is tethered to the sample chamber surface via binding between traptavidin and 2xbiotin. The chamber surface was passivated by molecular cushion of PEG polymers. The dashed lines on the top represent magnetic fields generated by a pair of permanent magnets. Lipids and detergents of the bicelle are denoted in blue and green, respectively. Figure 1–figure supplement 1. Preparation of molecular samples. Figure 1–source data 1. Original files of raw unedited gels and figures with uncropped gels.

**Figure 2.**
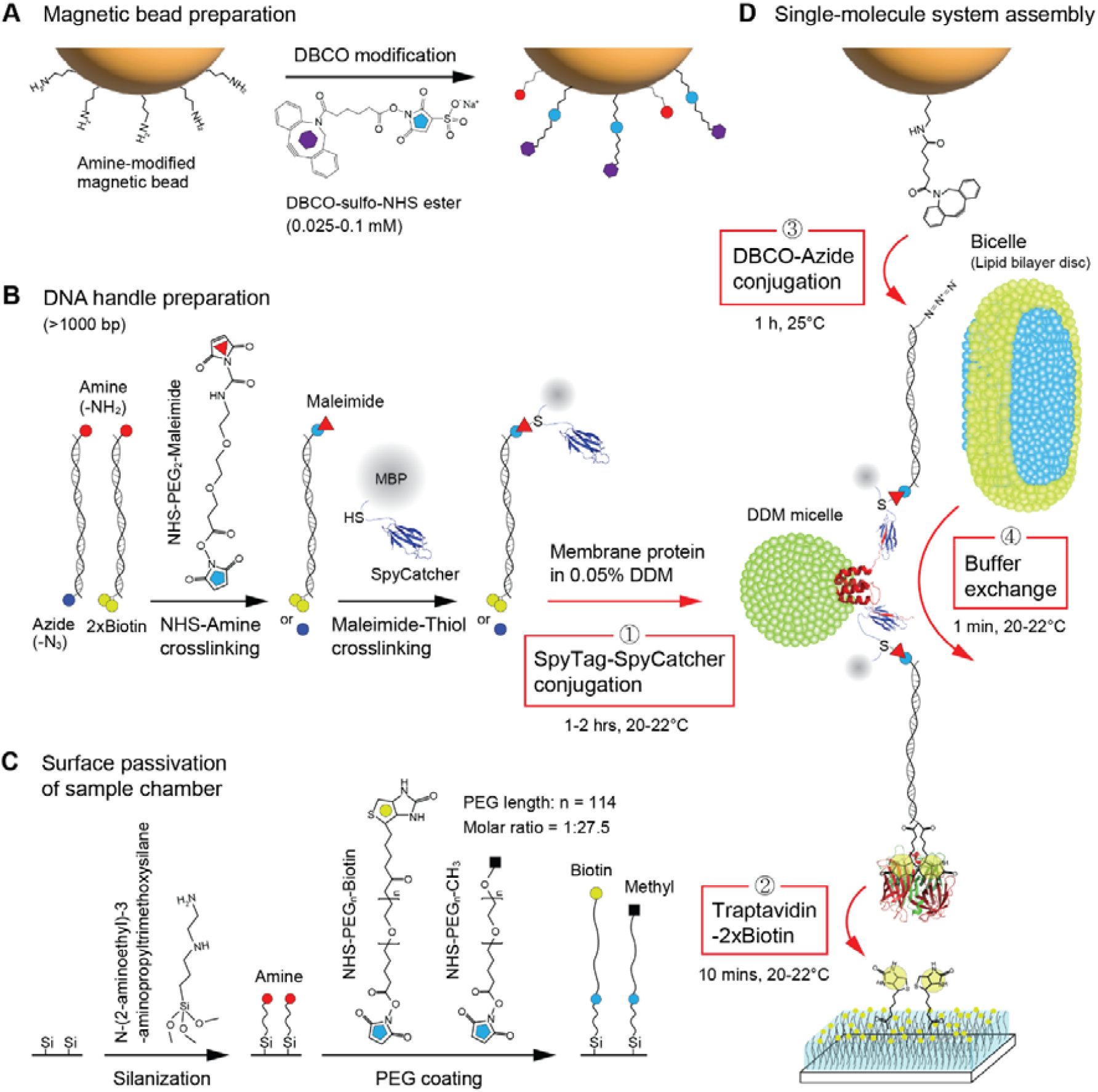
Procedure of single membrane protein tethering for mechanical manipulation. (A) DBCO modification on the surface of amine-coated magnetic beads using the DBCO-sulfo-NHS ester crosslinker (optimal conc. = 0.025-0.1 mM). (B) Preparation of SpyCatcher-DNA handles via thiol-maleimide conjugation. One end of the handles (>1000 bp) is labeled with amine, while the other end of the handles is labeled with azide or 2xbiotin. The amine end is modified by maleimide using the NHS-PEG_2_-Maleimide crosslinker, followed by the attachment to SpyCather fused with maltose binding protein (MBP) at the cystine residue. (C) Passivation of sample chamber surface using PEG polymers. The chamber surface is functionalized by amines via silanization, followed by PEG coating via NHS-amine conjugation. For the surface passivation, a high concentration of biotin-PEG was added to methyl-PEG solution at a 1:27.5 molar ratio. (D) Assembly of single-molecule pulling system. The SpyCatcher-DNA handles are attached to a membrane protein solubilized in 0.05% DDM via covalent SpyTag/Cather conjugation (1^st^ step). The hybrid molecular construct is bound to traptavidin at its 2xbiotin-modified end. The remaining biotin-binding pockets of the traptativin are then attached to biotins highly passivated on the chamber surface (2^nd^ step). The DBCO-modified magnetic bead is tethered to the other end of the molecular construct via DBCO-azide conjugation (3^rd^ step). The final buffer exchange with bicelles allows for lipid bilayer environment for the singly-tethered membrane protein (4^th^ step).

For the DNA handle attachment, we used the SpyTag/SpyCatcher conjugation system^35,37^ (Figure 2B and Figure 1–figure supplement 1). Amine-labeled end of the DNA handles was modified to maleimide using a crosslinker with the reactive groups of N-hydroxysuccinimide (NHS) ester and maleimide, and then attached to the thiol of a cysteine engineered in SpyCatcher (see Methods). The SpyCatcher-labeled DNA handles were then bound to the SpyTags, a short 13-amino-acid peptide tag, at the protein ends (Figure 2D; 1-2 hrs at 20-22°C). Upon rapid binding, an isopeptide bond between the pair spontaneously forms^37^. The other end of the handles was modified with azide or 2xbiotin to tether the molecular constructs to the magnetic beads and sample chamber surface, respectively (Figs. 1,2). The DNA hairpin with azide modification at one end was conjugated to a 2070-bp long DNA handle with 2xbiotin modification via annealing and ligation (Figure 1; see Methods).

The tetrameric traptavidin with four biotin-binding pockets was attached to the 2xbiotin-modified DNA handle (Figure 2D). The traptavidin is a streptavidin variant with 17-fold slower biotin dissociation (off-rate constant ratio: *k*_off,S_/*k*_off,T_ = 16.7) and 8-fold stronger binding (dissociation constant ratio: *K*_d,S_/*K*_d,T_ = 8.3)^38^. The remaining biotin-binding pockets of the traptavidin-attached molecular constructs were then tethered to the sample chamber surface passivated by a high concentration of biotinylated polyethylene glycol (biotin-PEG) among methyl PEG (mPEG) (∼1 biotin-PEG per 5 × 5 mPEG molecules) (Figure 2D; 10 mins at 20-22°C). The high concentration and long polymer arm (∼32 nm contour length) of biotin-PEG likely allow for the capture of all remaining binding pockets of traptavidins^39^. The polymer cushion of mPEG prevents nonspecific adhesion of the sticky DBCO-coated beads and highly hydrophobic molecules, such as membrane proteins (Figure 2C).

To tether the other end of the molecular constructs to the magnetic beads, we employed the click conjugation between DBCO and azide moieties, where the azide-modified end of the molecular constructs was conjugated to the DBCO-modified magnetic beads without catalytic enzymes or metal ions, simplifying the system (Figure 2A). DBCO modification was performed on amine-functionalized beads using a DBCO-NHS-ester crosslinker (see Methods). The DBCO modification on the bead surface was a critical step (optimal conc. = 0.025-0.1 mM) because lower or higher DBCO concentrations (*e*.*g*., 0.01 or 1 mM) resulted in unsuccessful conjugation to target molecules or entire nonspecific binding of beads to the surface, respectively (Figure 3 and Figure 3–source data 1).

**Figure 3.**
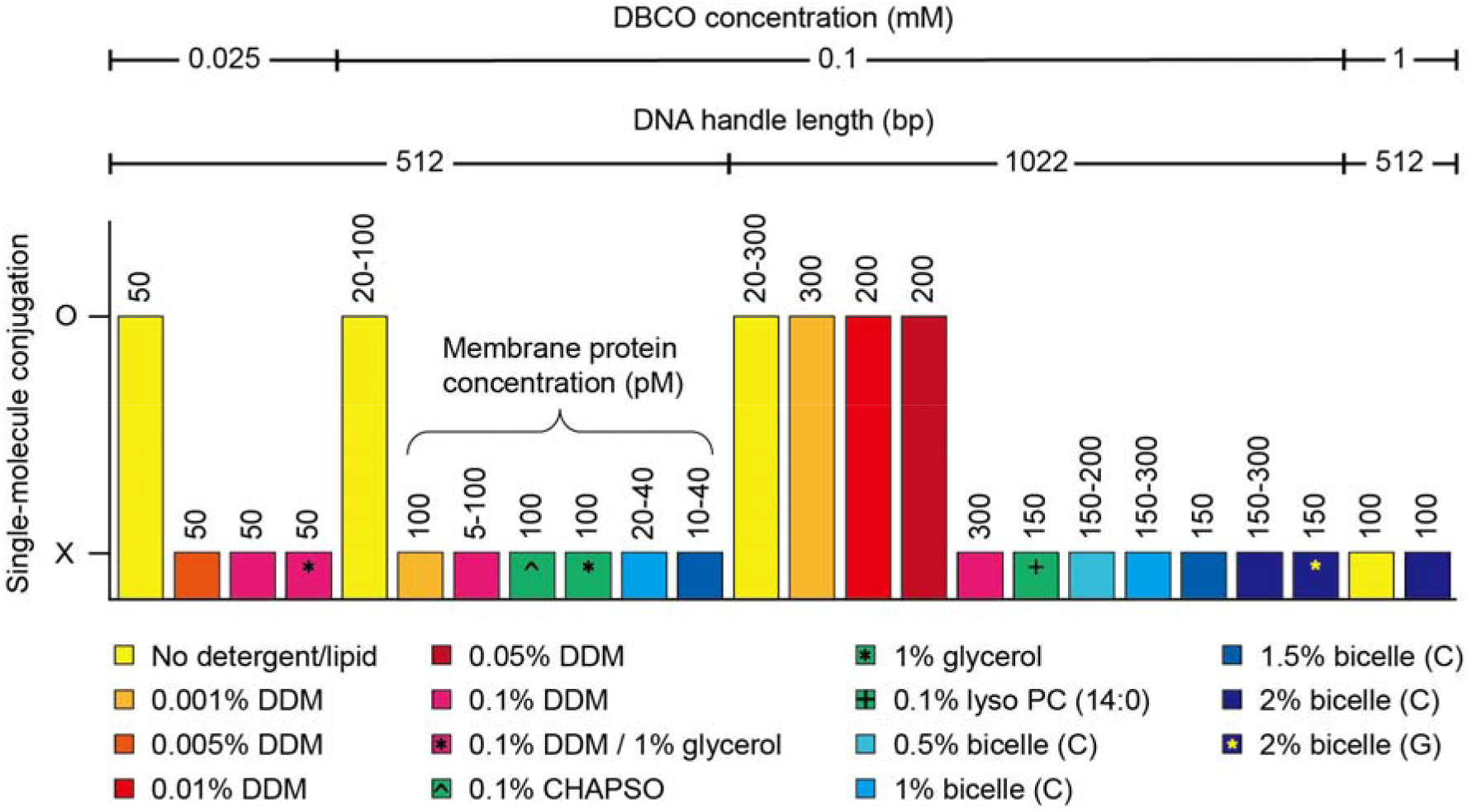
Searching conditions for successful DBCO-azide conjugation in single-molecule tethering. The DBCO concentration indicates the final concentration of the DBCO-sulfo-NHS ester crosslinker in the reaction of DBCO modification on bead surface. Others, such as DNA handle length, membrane protein concentration, and detergent concentration, indicate the conditions in the DBCO-azide conjugation reaction in single-molecule sample chambers. The base buffers are a phosphate-buffered saline (100 mM sodium phosphate, 150 mM NaCl, (pH 7.3) and a tris-buffered saline (50 mM Tris, 150 mM NaCl, pH 7.5). The bicelle (C or G) is composed of DMPC (or DMPG) lipid and CHAPSO detergent at a 2.5:1 molar ratio. The % indicates w/v %. The symbol O indicates successful conjugation between DBCO-modified magnetic beads and target membrane proteins so that specifically-tethered beads can be found and force-extension curve data can be obtained (confirmed by three replicates). The symbol X indicates unsuccessful conjugation to target proteins or entire nonspecific binding of beads to the chamber surface. In this case, no data is obtained. See Figure 3–source data 1 for more details. Figure 3–source data 1. Full list of tested conditions for DBCO-azide conjugation.

Membrane proteins require a suitable solubilization environment, such as bicelles or detergent micelles. Determining the optimal conditions for solubilization during the DBCO-azide conjugation reaction was another critical step since the conjugation efficiency decreased presumably due to the nonpolar character of the lipids and detergents. Among tested conditions of various concentrations of DDM, DMPC-bicelle, DMPG-bicelle, CHAPSO, lyso PC, and glycerol (Figure 3 and Figure 3–source data 1), we found that 0.05% (w/v) DDM detergent enabled the bead tethering via DBCO-azide conjugation, as well as the stable solubilization of membrane proteins under micelle conditions (critical micelle concentration of DDM = ∼0.01%).

The length of the DNA handles was also critical in the tethering of DBCO-coated beads to the azide-modified handle of molecular constructs. Shorter handles of ∼500 bp or less that were used for single-molecule tweezers^10,11,17,19,20^ were not able to tether to the beads under the conditions of bicelles or detergent micelles (Figure 3 and Figure 3 –source data 1). This result is perhaps due to the reduced radius-of-gyration of shorter handles confined to the chamber surface, reducing the efficiency of the DBCO-azide conjugation in bicelle/detergent solutions. After incubation of beads in the single-molecule chamber (1 h at 25°C), unconjugated beads were washed. The detergent micelles were then exchanged with bicelles to reconstitute the lipid bilayer environment for membrane proteins (Figure 2D; 1 min at 20-22°C).

The single-molecule magnetic tweezers using the strong tethers allowed for repetitive mechanical pulling on the membrane protein, scTMHC2, or the DNA hairpin, 17S6L, in force scanning of 1-30 pN (Figure 4A,B). scTMHC2 was unfolded at 18.2 ± 0.1 pN (*N* = 5 molecules, *n* = 1000 data points; Figure 4–figure supplement 1), consistent with a previous study^34^. The scatter plot of unfolding forces and step sizes of scTMHC2 appeared in the helix-coil transition zone bounded by force-extension curves for its unfolded polypeptide state (*U*_p_) and unfolded helical state (*U*_h_) (Figure 4C; *N* = 5 molecules, *n* = 1000 data points). The force-extension curves of the two extreme states were modeled by worm-like chain and Kessler-Rabin polymer models (see Methods). The area of the scatter plot data points around 18 pN matched that of the previously-reported helix-coil transition of a helical membrane protein, GlpG^33,36^.

**Figure 4.**
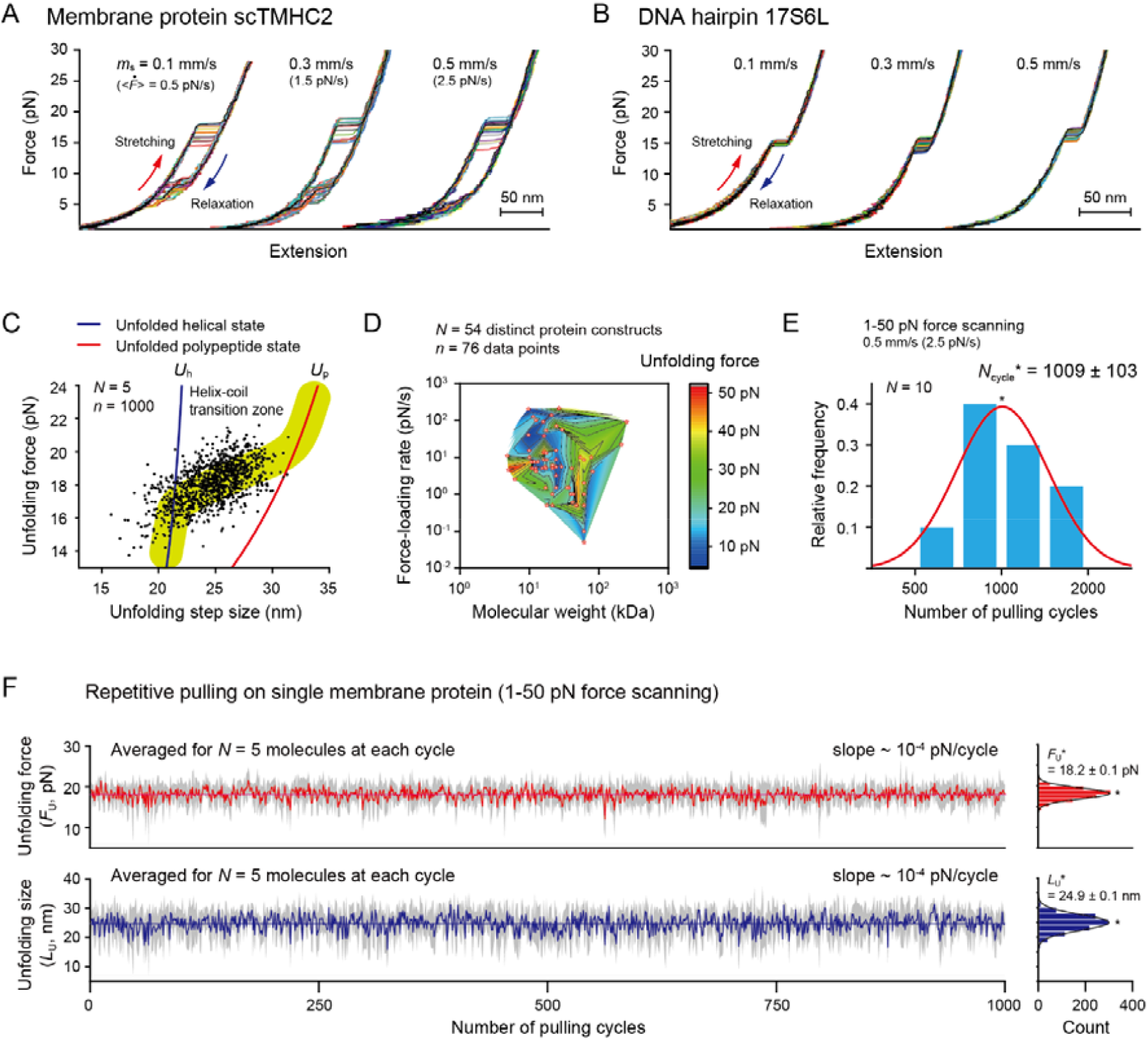
Evaluation of robustness of our single-molecule tweezers. (A,B) Representative force-extension curves of the membrane protein scTMHC2 (A) and the DNA hairpin 17S6L (B) at different pulling speeds (*N* = 20 for each condition). The pulling speed (magnet speed) is denoted as *m*_s_. The average force-loading rate in 1-50 pN is denoted as <*F* >. (C) Scatter plot of unfolding forces and step sizes of the membrane protein (*N* = 5 molecules, *n* = 1000 data points). The blue and red lines represent the force-extension curves for its unfolded helical state (*U*_h_) and unfolded polypeptide state (*U*_p_), respectively. (D) Survey of published protein unfolding forces that were experimentally measured (*N* = 54 distinct protein constructs, *n* = 76 data points; see Figure 4–source data 1). 98% of them is less than 50 pN that is set as the practical upper limit of the force. (E) Relative frequency of the number of pulling on single membrane protein (*N* = 10 molecules). The peak value of gaussian-fit function (*N*_cycle_*) is 1009 ± 103 (standard error, SE). The standard deviation (SD) of the distribution is 530. The force-scanning range was 1-50 pN. The pulling speed and average force-loading rate were 0.5 mm/s and 2.5 pN/s, respectively. (F) Unfolding force and step size as a function of the number of pulling on single membrane protein (analyzed for 5 molecules surviving more than the thousand cycles). The traces show the stability of the measured values in repetitive pulling. The first-order polynomial fitting yielded ∼10^−4^ pN/cycle for the slope of the traces. The colored traces are averaged ones over 5 different molecules at each cycle. The SD at each cycle is denoted in grey. Count histograms of the averaged traces are shown on the right (peak value ± SE). The SDs of the distributions are 1.3 pN and 2.7 nm, respectively. Figure 4–figure supplement 1. Distributions of unfolding forces and step sizes of the membrane protein scTMHC2. Figure 4–figure supplement 2. Unfolding and refolding forces as a function of the number of pulling on the DNA hairpin 17S6L. Figure 4–source data 1. Survey of protein unfolding forces in optical and magnetic tweezers. Figure 4–source data 2. Raw data for the figure panels.

To evaluate the robustness of the tweezers, we set a practical upper limit of the force that can be applied to most biological processes. For 98% of the 54 distinct protein constructs studied by optical/magnetic tweezers, the most probable or average unfolding forces were less than 50 pN, regardless of their molecular weights and force-loading rates (Figure 4D and Figure 4–source data 1). Thus, an arbitrary value of 50 pN was set as the upper force limit for each pulling cycle. The protein fold represents one of the strongest biomolecular interactions involving hundreds of amino acid residues. Therefore, most biomolecular interactions are testable below the 50 pN force limit.

With force scanning of 1-50 pN, our single-molecule tweezers were able to perform, on average, 1009 ± 103 cycles of repetitive mechanical manipulations on a single membrane protein until the dissociation of biotins from traptavidin (Figure 4E; *N* = 10 molecules). The unfolding forces and step sizes averaged for a few molecules at each cycle were almost constant (slope ∼ 10^−4^) within thermal fluctuations over a thousand of pulling cycles (Figure 4F; analyzed for 5 molecules surviving more than the thousand cycles; Figure 4–figure supplement 2 for the DNA hairpin). These results suggest that our high-throughput method can expedite membrane protein studies using single-molecule tweezers, which have been largely hindered by weak molecular tethering.

This approach of obtaining repetitive (un)folding or (un)binding of single molecules can determine the bond strength, kinetics, free energy landscape, and intermediate states^40,41^. The accuracy of the physicochemical quantities depends on the number of data points that are obtained from the single-molecule stretching. Thus, to evaluate the statistical reliability against the number of pulling cycles, we performed Monte Carlo (MC) simulations and compared the results with experiment (Figure 5; see Methods and Figure 5–figure supplements 1-4). We employed the unfolding kinetic parameters for this estimation: the unfolding rate constant (*k*_u0_) and the distance between the native and transition states (Δx_f_^†^), which can be extracted from the statistics of the unfolding force data. The kinetic values were obtained by analyzing the normalized plot of cumulative counts of unfolding forces (*i*.*e*., unfolding probability) using the rate equation and the Bell kinetic model^36^ (Figure 5–figure supplements 1-2; see Methods). The kinetic values, which were obtained from the experiment and simulations and then averaged for 5 molecules every 50 pulling cycles, presented the kinetic progress as a function of the number of pulling cycles (Figure 5A; Figure 5–figure supplement 3 for the DNA hairpin).

**Figure 5.**
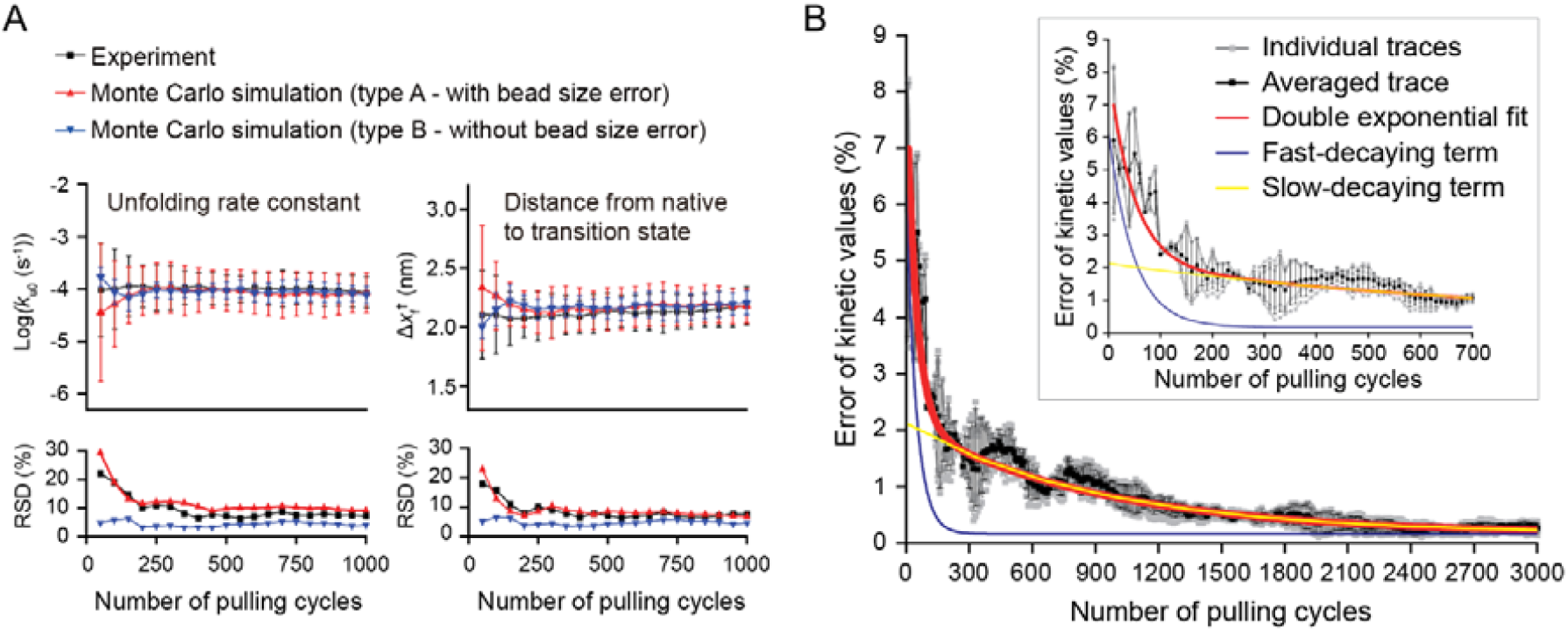
Evaluation of statistical reliability of our single-molecule tweezers. (A) Traces of kinetic parameters as a function of the number of pulling cycles that were obtained from the experiment and Monte Carlo (MC) simulations (*N* = 5 molecules). The *k*_u0_ and Δx_f_^†^ indicate the unfolding rate constant at zero force and the distance between the native and transition states, respectively. The relative standard deviation (RSD) indicates the ratio of the standard deviation to the mean. (B) Error of kinetic values as a function of the number of pulling cycles. The individual traces (grey) were created from MC simulations shown in Figure 5– figure supplement 4. The averaged trace (black) is the one averaged for the individual traces at each pulling cycle. The red curve represents the function of double exponential fit. The blue and yellow curves represent the fast- and slow-decaying term, respectively. The inset shows the initial phase of hundreds of pulling cycles. Figure 5–figure supplement 1. Unfolding and refolding probabilities as a function of force for the membrane protein and DNA hairpin. Figure 5–figure supplement 2. Comparison of two modes of kinetic analyses. Figure 5–figure supplement 3. Unfolding and refolding kinetic traces for the DNA hairpin as a function of the number of pulling cycles. Figure 5–figure supplement 4. Evaluation of kinetic errors for various shapes of unfolding probability profiles. Figure 5–source data 1. Raw data for the figure panels. Figure 5–source data 2. Codes for the Monte Carlo simulations

The relative standard deviations (RSD; defined as the ratio of the standard deviation to the mean) for the experiment and MC simulation with bead size error (type A) were a few to tens of % higher than those of the simulation without bead size error (type B) (Figure 5A, lower; Figure 5–figure supplement 3 for the DNA hairpin). However, the average kinetic values converged after the initial hundreds of cycles (Figure 5A, upper), indicating that the kinetic values averaged for a few molecules effectively excluded the effect of heterogeneous bead size after the initial phase. Using MC simulations, we quantitatively evaluated the average error of the kinetic values against the number of pulling cycles (Figure 5B and Figure 5–figure supplement 4). We found that the kinetic error rapidly decayed to zero during the initial 200 pulling cycles, followed by slow decay (Figure 5B, inset). With the correction of intrinsic kinetic error in single-molecule force spectroscopy^42^, the average error of kinetics can be approximated as 1.8% for 200 cycles or 0.8% for 1000 cycles of pulling (Figure 5B), which can be stably performed in our robust single-molecule tweezers.

## Discussion

Single-molecule tweezers have recently been utilized as an emerging tool for membrane proteins studies. Very low level of their spring constants (∼10^−4^ pN/nm) enables constant low-force measurements^43^. This advantage has allowed for high-resolution observation of low-energy TM helix interaction of helical membrane proteins within lipid bilayers^10^. However, the methods rely on conventional noncovalent tethers such as dig-antidig, which is as very weak as *τ*_off_ = ∼1 sec at 25 pN (ref. ^24^). This has limited the stable performance of nanomechanical manipulation on a single membrane protein and reliable statistical analyses. Our robust single-molecule tweezers using the DBCO-azide click conjugation and the traptavidin-2xbiotin complex enables more than 1000 cycles of mechanical pulling on a single molecule. The upper force limit of 50 pN for testing the system robustness covers most biomolecular interactions, up to 98% of the surveyed protein folds. Thus, our method permits the stable stretching experiments on diverse proteins in relevant force ranges, which will accelerate membrane protein studies using the tweezer approaches. The robust linkage system could also allow for mechanical manipulation studies under harsh conditions such as an environment of abundant reactive oxygen species.

Our single-molecule tweezers, which can perform a large number of mechanical manipulations at different time points over tens of hours, could address new questions on membrane proteins. For example, oxidative modification could be prominent for some membrane proteins as the time passes. The aging effect may alter the folding and function of membrane proteins. Indeed, single-molecule tweezers have revealed the phenomenon of the protein aging or folding fatigue of several water-soluble proteins, such as protein L, ubiquitin, and titin^44,45^. Thus, our tweezer method can be used to quantify the long-period evolution of biophysical properties of membrane proteins in more physiological, low-force ranges of a few to tens of pN. How chaperone proteins or small molecules like ligands act on the time-evolved, different protein populations can also be captured.

The results of the MC simulations show that, for proteins without temporal changes in an observed time span, our method can obtain statistically-reliable data points, such as for the unfolding kinetics within an approximate 2% error for 200 cycles (fast-decaying for the error). After that, 5 times larger number of pulling cycles up to 1000 does not sufficiently improve the accuracy, reducing the error to an approximate 1% (slow-decaying for the error). This indicates that blindly applying a large number of stretching cycles on single molecules is not profitable in a practical sense. Thus, it should be beneficial to set practical limits of pulling cycles by examining the error-decay profiles for measured quantities.

The single-molecule tethering strategies described here are not restricted to membrane proteins (as shown for the DNA hairpin) or specific tweezer apparatus. One advantage of our method is that the covalent conjugation using DBCO is specific to the azide moiety and does not require catalytic enzymes or metal ions, simplifying the system. Thus, the DBCO-mediated tethering can be considered as an additional and orthogonal strategy for single protein manipulation. Another advantage is that the molecular tether using traptavidin (a streptavidin variant with 17-fold slower biotin dissociation and 8-fold stronger binding) is one of the strongest and rapid noncovalent tethers, which can be widely utilized in single-molecule tweezers. A third advantage is that pulling a single protein instead of a tandem repeat of proteins, with hundreds-of-nm DNA handles attached via the facile SpyTag/Catcher conjugation, excludes the possibility of undesirable interactions and aggregation between proteins. It also allows for native protein studies without cysteine mutations. We anticipate that all tethering strategies established here can be widely adopted in other versions of magnetic tweezers, optical tweezers, and their hybrid apparatuses with fluorescence imaging, to improve their system stability.

## Methods

### Traptavidin purification

Traptavidin with 6xHis tag at the C-terminus encoded in pET24a vector was transformed into *E. coli* BL21-Gold(DE3)pLysS (Agilent). A selected colony from a transformed agar plate was grown in 1 L of Luria-Bertani (LB) medium with 25 mg/ml kanamycin at 37°C. At approximately OD_600_ = 0.9, 0.5 mM IPTG was added to overexpress the protein for 4 hrs at 37°C. The cell culture was centrifuged at 5700 rpm for 10 mins at 4°C and the cell pellet was resuspended in 50 ml lysis buffer (10 mM Na_2_HPO_4_, 2 mM KH_2_PO_4_, 287 mM NaCl, 2.7 mM KCl, 5 mM EDTA, 1% Triton X-100, pH 7.4). The resuspension was lysed with Emulsiflex C3 high pressure homogenizer (∼17000 psi, Avestin). The cell lysate was centrifuged at 18000 rpm for 30 mins at 4°C, the supernatant was removed, and the inclusion body pellet was washed with 10 ml extraction buffer (10 mM Na_2_HPO_4_, 2 mM KH_2_PO_4_, 137 mM NaCl, 2.7 mM KCl, 0.5% Triton X-100, pH 7.4). The washing step was repeated three times. The washed pellet was dissolved in 6 M guanidine hydrochloride (GuHCl), followed by dilution into refolding buffer (10 mM Na_2_HPO_4_, 2 mM KH_2_PO_4_, 137 mM NaCl, 2.7 mM KCl, 10 mM imidazole, pH 7.4) and then incubation for 16 hrs at 4°C (refs. ^38,46^). The sample was centrifuged at 17700 g for 15 mins at 4°C. The supernatant was incubated with 2 ml Ni-IDA resin (Takara Bio) for 2 hrs at 4°C, and then washed in a gravity column with 10 ml wash buffer (10 mM Na_2_HPO_4_, 2 mM KH_2_PO_4_, 287 mM NaCl, 2.7 mM KCl, 30 mM imidazole, pH 7.4) three times. The protein sample was eluted with elution buffer (10 mM Na_2_HPO_4_, 2 mM KH_2_PO_4_, 287 mM NaCl, 2.7 mM KCl, 300 mM imidazole, pH 7.4), purified by size exclusion chromatography (HiLoad 16/600 Supderdex 75pg, Cytiva), and stored at -80°C in aliquots (Figure 1–figure supplement 1).

### Membrane protein purification

The membrane protein scTMHC2 adopted for this study is a designed single-chain TM homodimer that was previously reported^34^. The SpyTag sequence with a linker GGSGGS at its N- and C-termini was encoded for the DNA handle attachment. The gene block was cloned into pBT7-C-His vector (Bioneer) and the vector was inserted into *E. coli* BL21-Gold(DE3)pLysS by heat shock transformation (40 sec at 42°C). The cells were grown in LB broth with 100 μg/ml ampicillin at 37°C until OD_600_ reached to approximately 1.0. The overexpression was induced by the addition of 200 μM IPTG. The protein was expressed for 18 hrs at 18°C and the cells were harvested by centrifugation (5700 rpm, 10 mins, 4°C). The cell pellets were resuspended in 50 ml of a lysis buffer (25 mM Tris-HCl, 150 mM NaCl, 10% glycerol, 1 mM TCEP, 1 mM PMSF, pH 7.4). The sample was lysed by Emulsiflex-C3 at the pressure of 15000-17000 psi. 1% of DDM detergent (GoldBio) was added to the lysed cell and the cells was incubated for 1 h at 4°C to extract the membrane protein. The cell debris was removed by centrifugation (18,000 rpm, 30 mins, 4°C) and the supernatant was incubated with Ni-IDA resin for 1 h at 4°C. The resin was packed into a column and was washed with a washing buffer (25 mM Tris-HCl, 150 mM NaCl, 10% glycerol, 1 mM TCEP, 0.1% DDM, 20 mM imidazole, pH 7.4). The protein was then eluted with elution buffer (25 mM Tris-HCl, 150 mM NaCl, 10% glycerol, 1 mM TCEP, 0.1% DDM, 300 mM imidazole, pH 7.4). The purified membrane protein was concentrated to ∼10 µM and stored at -80°C in aliquots (Figure 1–figure supplement 1).

### DNA handle attachment to membrane proteins

Two types of 1022-bp DNA handles (modified with amine at one end; azide or 2xbiotin at the other end) were amplified by PCR from λ DNA template (NEB, N3011S). The primers were added to total 8 ml PCR mixture: 1 µM of a forward primer (ACAGAAAGCCGCAGAGCA) with amine at 5’ end, 0.5 µM of a reverse primer (TCGCCACCATCATTTCCA) with azide at 5’ end, and 0.5 µM of the reverse primer with 2xbiotin at 5’ end. The DNA handles were purified using HiSpeed Plasmid Maxi kit (Qiagen). For maleimide modification at the amine end, the purified DNA in 1 ml NaHCO_3_ (pH 8.3) was incubated with 1 mM SM(PEG)_2_ (Thermo Scientific Pierce) for 20 mins at 20-22°C. Unconjugated SM(PEG)_2_ was removed by Econo-Pac 10DG Column (Bio-Rad). The buffer used for column equilibration and sample elution was 0.1 M sodium phosphate (pH 7.3) with 150 mM NaCl. The DNA handles modified with maleimide were covalently attached to MBP-fused SpyCatcher with the single cysteine (2 hrs at 20-22°C for the incubation; ∼20 µM for the protein concentration)^35^. Unconjugated proteins were removed by HiTrap Q HP column (Cytiva) with the gradient mode of 0-1 M NaCl in 20 mM Tris-HCl (pH 7.5). The DNA peak fractions were concentrated to ∼350 nM and stored at -80°C in 10 μl aliquots. Approximately 30% of the sample is the SpyCather-conjugated DNA handle (∼100 nM). The conjugated DNA handles were attached to the SpyTagged target proteins (1-2 hrs at 20-22°C for the incubation; ∼70 nM for DNA and ∼2 µM for protein). The sample was diluted to make ∼200 pM DNA handle and stored at -80°C in 10 μl aliquots (Figure 1–figure supplement 1).

### Making DNA hairpin construct

A 2070-bp DNA handle was amplified by PCR from λ DNA template (final 2.5 µg/ml; NEB, N3011S) using a forward primer (TAAGGATGAACAGTTCTGGC) with 2xbiotin at 5’ end, and a reverse primer (GCAGCGAGTTGTTCC/1’,2’-Dideoxyribose/AATGATCCATTAATGG CTTGG) (final 2 µM each). The total 200 µl PCR product was purified with HiGene™ Gel & PCR Purification System (Biofact, GP104-100) with 50 µl elution buffer (10 mM Tris-HCl, pH 8.0). The purified sample was diluted to 100 nM with deionized water. 50 nM of DNA handle was mixed with 500 nM of an 80-nt single-stranded DNA forming a hairpin structure (GAACAACTCGCTGCAAACAACAACAAAAGAGTCAACGTCTGGATCTTTTTTGATCCAGACGTTGACTCAAAAGACATACC). The hairpin DNA has phosphate at 5’ end and azide at 3’ end. The incubation condition was 1 h at 37°C in ligation buffer (50 mM Tris-HCl, 10 mM MgCl_2_, 10 mM DTT, 10 mM ATP, pH 7.5). The mixture was further incubated for 16 hrs at 16°C with T4 DNA ligase added (Enzynomics, M001S). After the ligation, the ligase was inactivated by the sample incubation at 65°C for 10 mins followed by 5 mins on ice. The sample was purified with the purification kit with 50 µl of the elution buffer. The ligation yield was approximately 30%. The ligated product was diluted to 1 nM with 10 mM Tris-HCl (pH 8.0) with 0.1 mM EDTA, and stored at -20°C in 10 µl aliquots.

### DBCO modification on magnetic bead surface

The surface of magnetic beads was modified by DBCO using the conjugation between amine and NHS ester. 35 µl of amine-labeled magnetic beads (Thermo Fisher Scientific, M-270 amine, 14307D) was washed with buffer A (0.1 M sodium phosphate, 150 mM NaCl, 0.01% Tween20, pH 7.4) using DynaMag™-2 Magnet (Thermo Fisher Scientific, 12321D). After the wash, the beads were resuspended with 100 μl of the buffer A. The bead slurry was incubated with 10 µl of 0.25-1 mM DBCO-sulfo-NHS (Sigma, 762040; dissolved in DMSO) for 3 hrs at 25°C with slow rotation. The bead slurry was washed again and resuspended in 35 µl of the buffer A. The DBCO-modified bead sample was stored at 4°C.

### Surface passivation of single-molecule sample chamber

The coverslips of 24×50 mm and 24×40 mm (VWR; No. 1.5) were used for the bottom and top surface of single-molecule sample chamber, respectively. Both coverslips were cleaned by 1 M KOH in sonication bath for 30 mins and washed with deionized water. The bottom coverslips (24×50 mm) were further cleaned by Piranha solution (H_2_SO_4_(98%):H_2_O_2_(30%) = 2:1 volume ratio). The surface of the coverslips was functionalized by amine using the silanization solution of N-(2-aminoethyl)-3-aminopropyltrimethoxysilane (Sigma-Aldrich, 8191720100), acetic acid, and methanol (1:5:100 volume ratio)^35^. To this end, the coverslips were incubated with the silanization solution for 30 mins at 20-22°C. The amine-functionalized coverslips were washed with methanol, deionized water, and dried with centrifugation (407 g, 5 mins). The PEG polymers (Laysan Bio; 1:27.5 molar ratio of Biotin-PEG-SVA-5000 and MPEG-SVA-5000) were conjugated to the surface of the bottom coverslips using the amine-NHS conjugation. To this end, each 50 μl of PEG solution (total ∼40 mM PEG mixture in 100 mM sodium bicarbonate, pH 8.4) was incubated between two coverslips for 4 hrs at 20-22°C in humidity chambers. The coverslips were washed with deionized water, dried with centrifugation (407 g, 5 mins), and stored at -20°C.

### Single-molecule tweezer experiments

The single-molecule experiments were performed on a custom-built magnetic tweezer apparatus as previously described^19^. The sample chamber was constructed with the two surface-treated coverslips with double-sided tapes. Its channel volume (1 CV) was ∼10 µl. Streptavidin-coated polystyrene beads (Spherotech, SVP-10-5) was used as reference beads for the stage drift correction. They were washed with buffer B (0.1 M sodium phosphate, 150 mM NaCl, 0.1% Tween 20, pH 7.4), injected into the chamber, and then incubated for 2-5 mins at 20-22°C. After incubation of 100 mg/ml bovine serum albumin (BSA) for 5 mins at 20-22°C, the chamber was washed with washing buffer I (for the membrane protein, 50 mM Tris-HCl, 150 mM NaCl, 0.1% DDM, pH 7.4; for the DNA hairpin, 0.1 M sodium phosphate, 150 mM NaCl, pH 7.4). The sample of target biomolecules (the membrane protein or DNA hairpin) was bound to traptavidins by incubating 10 µl of ∼200 pM sample and 1 µl of 0.04 µM traptavidin for 15 mins at 20-22°C. 1 CV of the sample (final 40∼200 pM) was injected into the sample chamber and incubated for 10 mins at 20-22°C. To block unoccupied biotin-binding pockets, 1 CV of a 30-nt biotin-labeled oligonucleotide (10 µM in the washing buffer I) was injected into the chamber and incubated for 5 mins at 20-22°C. The chamber was then washed with washing buffer II (50 mM Tris-HCl, 150 mM NaCl, 0.05% DDM, pH 7.4) for the membrane protein or washing buffer I for the DNA hairpin. The DBCO-modified magnetic beads were washed and resuspended with washing buffer II for the membrane protein and washing buffer I for the DNA hairpin (20x diluted). The magnetic beads were injected into the chamber and incubated for 1 h at 25°C. The chamber was washed with single-molecule buffer (for the membrane protein, 50 mM Tris-HCl, 150 mM NaCl, 2% bicelle, pH 7.4; for the DNA hairpin, 0.1 M sodium phosphate, 150 mM NaCl, pH 7.4). The bicelle is composed of DMPC lipid (Avanti) and CHAPSO detergent (Sigma) at a 2.5:1 molar ratio. The tweezer rooms are maintained at 20-22°C. Only specifically-tethered molecules are examined to obtain the force-extension spectra.

### Analysis of unfolding force and step size

Experimental data of the unfolding force and step size of scTMHC2 were compared with the force-extension curves for its unfolded polypeptide and helical states^33,36^. The force-extension curve of the unfolded polypeptide state (*U*_p_) without secondary helices was constructed by the worm-like chain model with the persistence length of 0.4 nm (refs. ^33,36,47^) and the contour length that was estimated by the number of unfolded residues of tertiary structure (*n* = 153 for scTMHC2) times the average residue-residue distance of 0.36 nm (refs. ^33,36,47^). For the unfolded helical state (*U*_h_) in which the TM helices are fully structured, the TM helices were modeled by the Kessler-Rabin model with the persistence length of 9.17 nm (refs. ^33,36^) and the contour length that was estimated by the number of helix residues times the average helical rise per residue of 0.16 nm (refs. ^33,36^). The polypeptide linkers in the unfolded helical state were modeled by the worm-like chain model. The force-extension curves were corrected by the end-to-end distance of the pulling residue points of tertiary structure (Δ*d* = 1.1 nm for scTMHC2) because the extension changes during unfolding are measured smaller by Δ*d*.

### Analysis of unfolding kinetics

Cumulative plots of the unfolding force distributions were normalized to the unfolding probabilities as a function of force. To obtain the values for the unfolding kinetic parameters (*k*_u0_ and Δ*x*_f_^†^), the unfolding probability profiles were fitted with the formula, 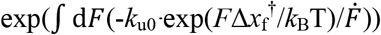, derived from the first-order rate equation, d*N*/d*t* = -*k*_u_*N*, and the Bell equation, *k*_u_ = *k*_u0_ exp(-*F*Δ*x*_f_^†^/*k*_B_T), where *F* is the force, *U* is the unfolded fraction (unfolding probability), *N* (=1-*U*) is the folded fraction (folding probability), *k*_u_ (*k*_u0_) is the kinetic rate constant at an arbitrary (zero) force, Δ*x*_f_^†^ is the distance between the native and transition states, *k*_B_ is the Boltzmann constant, *T* is the temperature, and 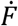 is the force-loading rate (d*F*/d*t*). The 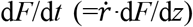 was approximated as a linear function of force (a*F*+b) in the force range of *U* = 0 to 1, where d*F*/d*z* is the first derivative of force with respect to magnet height that was calibrated by double exponential distribution and 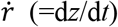 is the magnet speed that was maintained as constant in the single-molecule experiments. The constants of a and b were obtained by fitting the force-loading rate (d*F*/d*t*) as a function of force (*F*). To simplify the formula, *U* = 1-exp(d*F* (-*k*_u0_*·*exp(*F*Δ*x*_f_^†^/*k*_B_T)/(a*F*+b))), the exponential integral function (exp(*x*)/*x* d*x*) was approximated as the first term of series expansion (exp(*x*)/*x*) since higher terms are only ∼5% of the first term over the unfolding force ranges. The final equation used to fit the unfolding probabilities as a function of force was derived to *U* = 1-exp(-*k*_u0_*k*_B_*T·*exp(*F*Δ*x*_f_^†^/*k*_B_T)/(Δ*x*_f_^†^(a*F*+b))). The analysis yields the approximate values of unfolding kinetic parameters of *k*_u0_ and Δ*x*_f_^†^ (ref. ^36^).

### Monte Carlo simulations

In the Monte Carlo (MC) simulations (Figure 5 and Figure 5–figure supplements 1-4), unfolding forces were generated by random sampling from the probability density function (PDF) of unfolding as a function of force, *i*.*e*., the equation *U* = 1-exp(-*k*_u0_*k*_B_*T·*exp(*F*Δ*x*_f_^†^/*k*_B_T)/(Δ*x*_f_^†^(a*F*+b))) with set kinetic parameters, which was derived above. The cumulative force distributions were normalized to unfolding probabilities as a function of force. The unfolding probability profiles were then created in every 50 times of random sampling up to 1000-10000 times. Each unfolding probability was fitted with the equation to extract the kinetic values. The beads tethered to molecules have the error in their diameter (RSD < 3%; designated in the manual), which causes the error in unfolding forces and resultant probabilities. Thus, two types of MC simulations with or without bead heterogeneity were performed as simulation type A and B, respectively. For the type A, the random sampling was performed for all different PDFs that were experimentally measured, reflecting the error of bead size. For the type B, the random sampling was performed only for a median PDF with respect to the force at *U* = 0.5, effectively excluding the error of bead size.

## Acknowledgments

This work was supported and funded by the National Research Foundation of Korea (NRF-2020R1C1C1003937 to D.M.) and Ulsan National Institute of Science and Technology (1.190147.01 to D.M.). We thank S. Y. Hong for helpful comments.

## Author contributions

Seoyoon Kim: Methodology, Software, Validation, Formal analysis, Investigation, Resources, Data curation, Writing - Original Draft, Writing - Review & Editing, Visualization; Daehyo Lee: Investigation, Resources, Visualization; W.C. Bhashini Wijesinghe: Investigation, Resources; Duyoung Min: Conceptualization, Methodology, Software, Validation, Formal analysis, Investigation, Resources, Writing - Original Draft, Writing - Review & Editing, Visualization, Supervision, Project administration, Funding acquisition.

## Availability of data and codes

All data and analysis codes that support the findings of this study are available in the manuscript, figure supplements, source data, and source code files.

## Figure Supplements

Figure 1–figure supplement 1. Preparation of molecular samples.

Figure 4–figure supplement 1. Distributions of unfolding forces and step sizes of the membrane protein scTMHC2.

Figure 4–figure supplement 2. Unfolding and refolding forces as a function of the number of pulling on the DNA hairpin 17S6L.

Figure 5–figure supplement 1. Unfolding and refolding probabilities as a function of force for the membrane protein and DNA hairpin.

Figure 5–figure supplement 2. Comparison of two modes of kinetic analyses.

Figure 5–figure supplement 3. Unfolding and refolding kinetic traces for the DNA hairpin as a function of the number of pulling cycles.

Figure 5–figure supplement 4. Evaluation of kinetic errors for various shapes of unfolding probability profiles.

**Figure 1–figure supplement 1.**
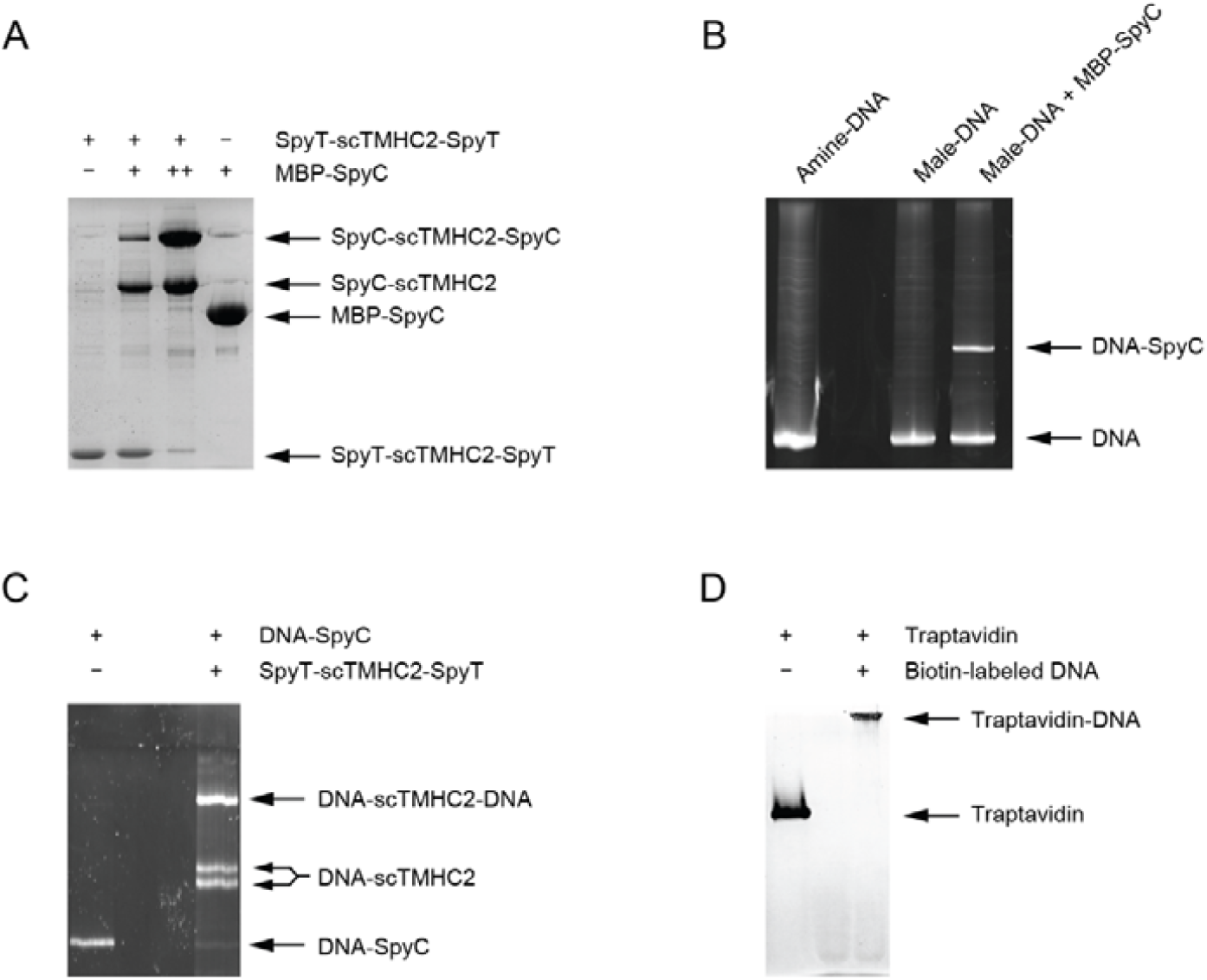
Preparation of molecular samples. (A) Purification of the membrane protein scTMHC2 confirmed by 10% SDS-PAGE. After protein purification, the binding test for SpyTag (SpyT) of scTMHC2 was performed by incubating with MBP-fused SpyCatcher (MBP-SpyC). MBP indicates maltose binding protein. Upon binding, the SpyT and SpyC spontaneously form an isopeptide bond. The doubly-attached construct become dominant when a higher concentration of MBP-SpyC (denoted as ++) is added. (B) MBP-SpyC attachment to DNA handles confirmed by 6% SDS-PAGE. The amine-labeled end of the DNA is modified to maleimide (male) using an NHS-male crosslinker, followed by the attachment to the cysteine thiol of MBP-SpyC. The yield is approximately 30%. (C) DNA handle attachment to scTMHC2 confirmed by 6% SDS-PAGE. The DNA handles conjugated to MBP-SpyC are attached to scTMHC2 via the binding between SpyT and SpyC. Upon binding, the SpyT and SpyC spontaneously form an isopeptide bond. The two bands for the singly-handled construct correspond to the N- and C-terminal attachments, respectively. Approximately 50% of the doubly-handled construct is the one with the modification of azide at one end and 2xbiotin at the other end. Only this construct can be tethered to magnetic beads and sample chamber surface as shown in Figure 1. (D) Purification of traptavidin confirmed by 12% SDS-PAGE. After protein purification, the binding test was performed by incubating with 100-bp biotinylated DNA.

**Figure 4–figure supplement 1.**
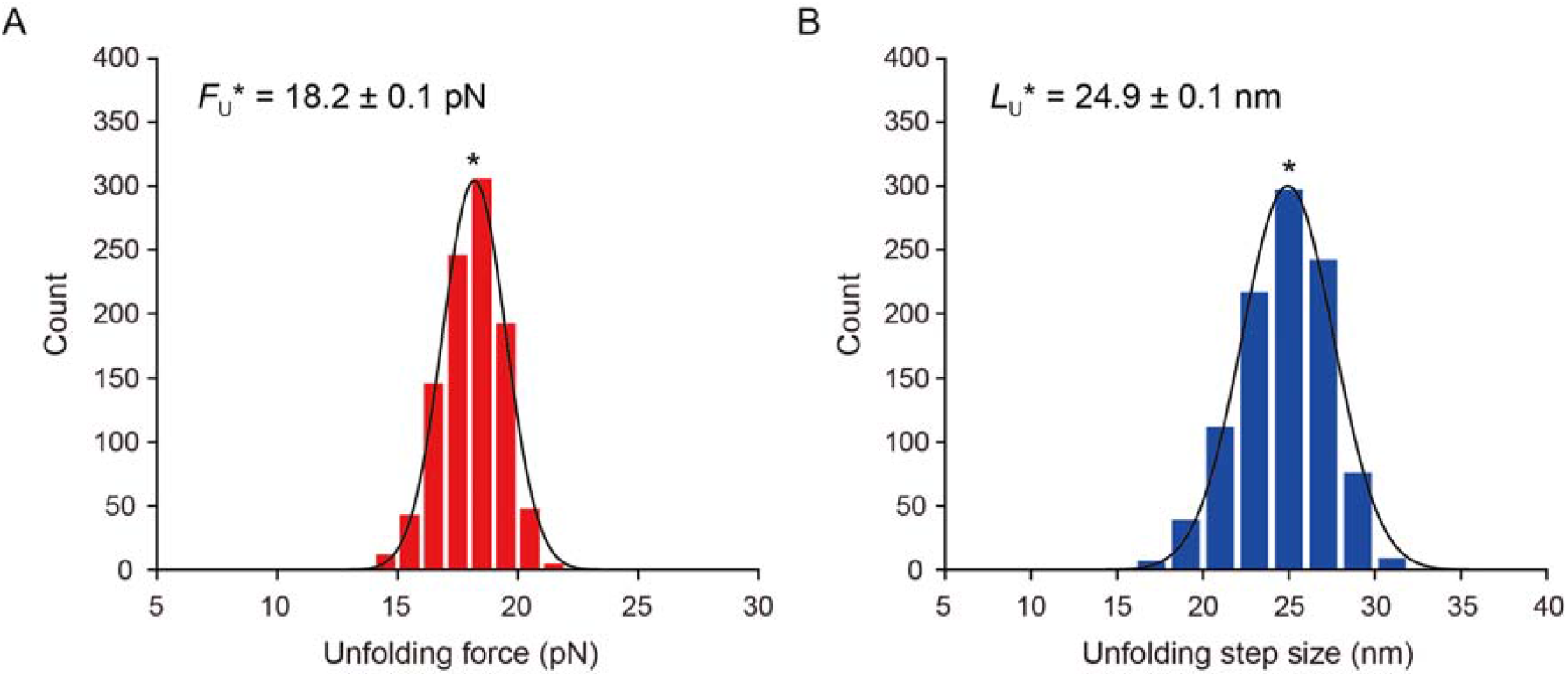
Distributions of unfolding forces and step sizes of the membrane protein scTMHC2. (A,B) Count histograms of unfolding forces (A) and step sizes (B). The count histograms were obtained from the averaged values at each pulling cycle as shown in Figure 4F (*N* = 5 molecules, *n* = 1000 data points). The gaussian fits yield most probable unfolding force and step size (peak value ± standard error, SE). The standard deviations of the distributions are 1.3 pN and 2.7 nm, respectively.

**Figure 4–figure supplement 2.**
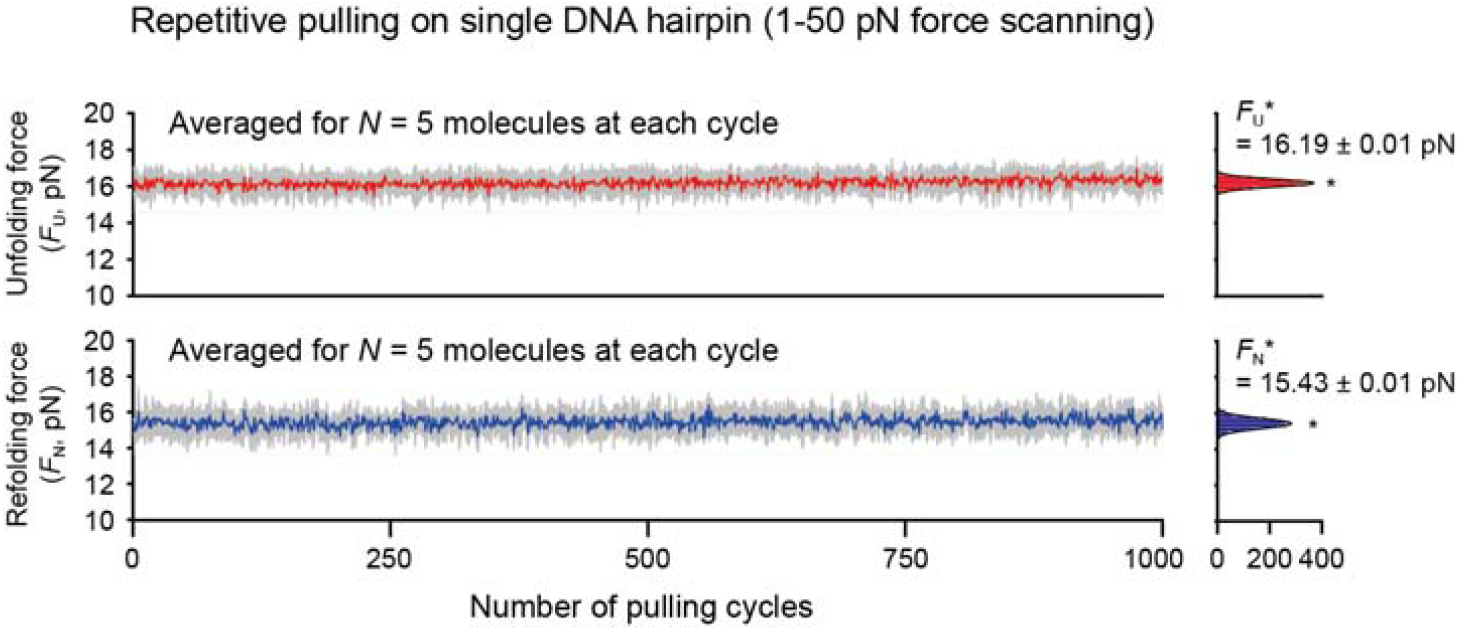
Unfolding and refolding forces as a function of the number of pulling on the DNA hairpin 17S6L. The data were analyzed for 5 molecules surviving more than 1000 pulling cycles. The traces show the stability of the measured values in repetitive pulling (constant within thermal fluctuations). The colored traces are averaged ones over 5 different molecules at each cycle. The standard deviation (SD) at each cycle is denoted in grey. Count histograms of the averaged traces are shown on the right (peak value ± SE). The SDs of the distributions are 0.21 pN and 0.28 pN, respectively.

**Figure 5–figure supplement 1.**
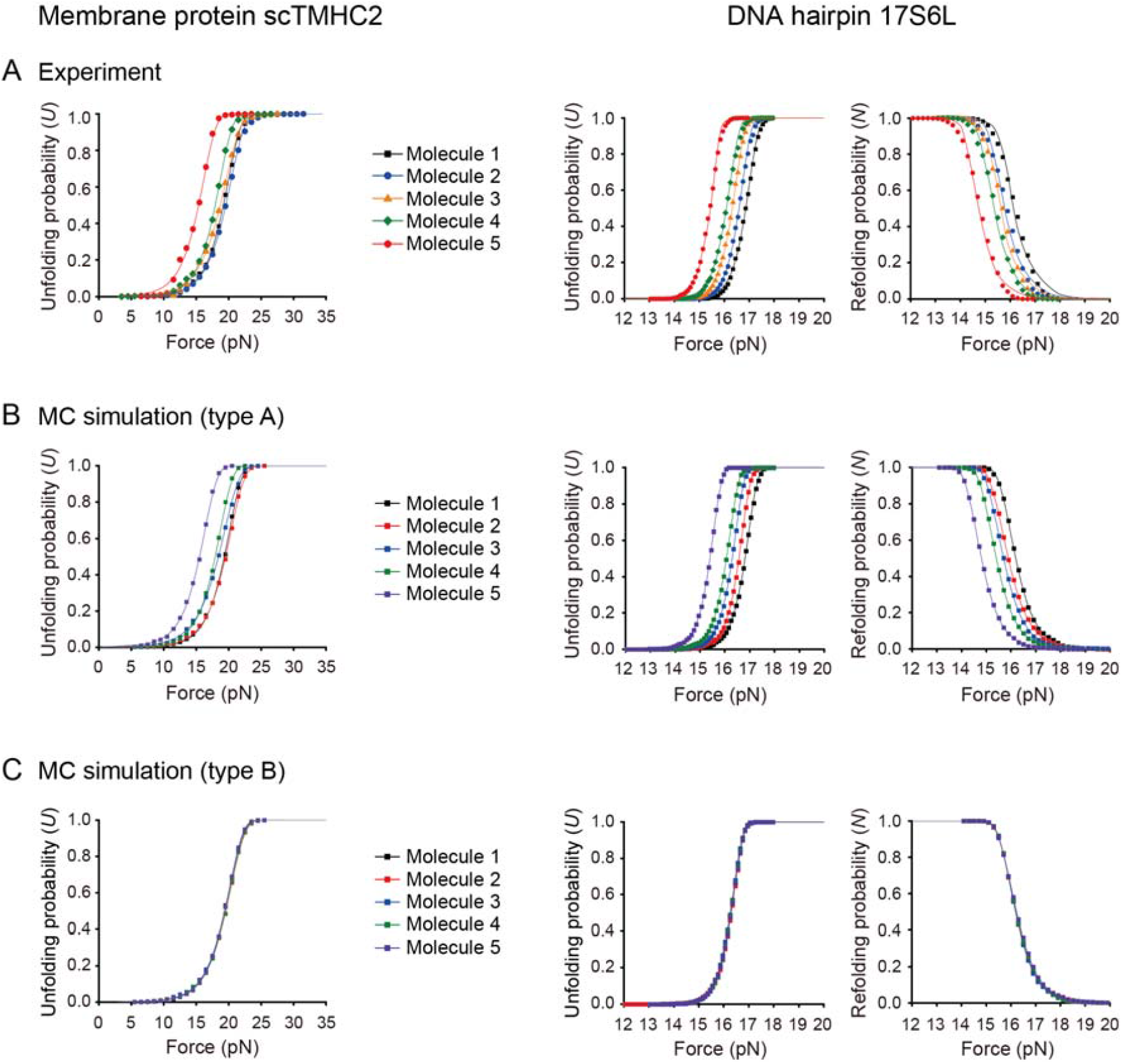
Unfolding and refolding probabilities as a function of force for the membrane protein and DNA hairpin. (A-C) Data obtained from the experiment (A) and Monte Carlo (MC) simulations (B,C) (*N* = 5 molecules; *n* = 1000 or 1500 data points for the membrane protein or DNA hairpin, respectively). In the type-A simulation, the force values of the unfolding and refolding were generated from all probability density functions (PDFs) obtained from the experiment, reflecting the error of bead size (see Methods). In the type-B simulation, the force values of the unfolding and refolding were generated from a median PDF with respect to the force at *U* = 0.5, effectively excluding the error of bead size. The data from different molecules in the type-B simulation indicate random sampling from the same PDF at different time points.

**Figure 5–figure supplement 2.**
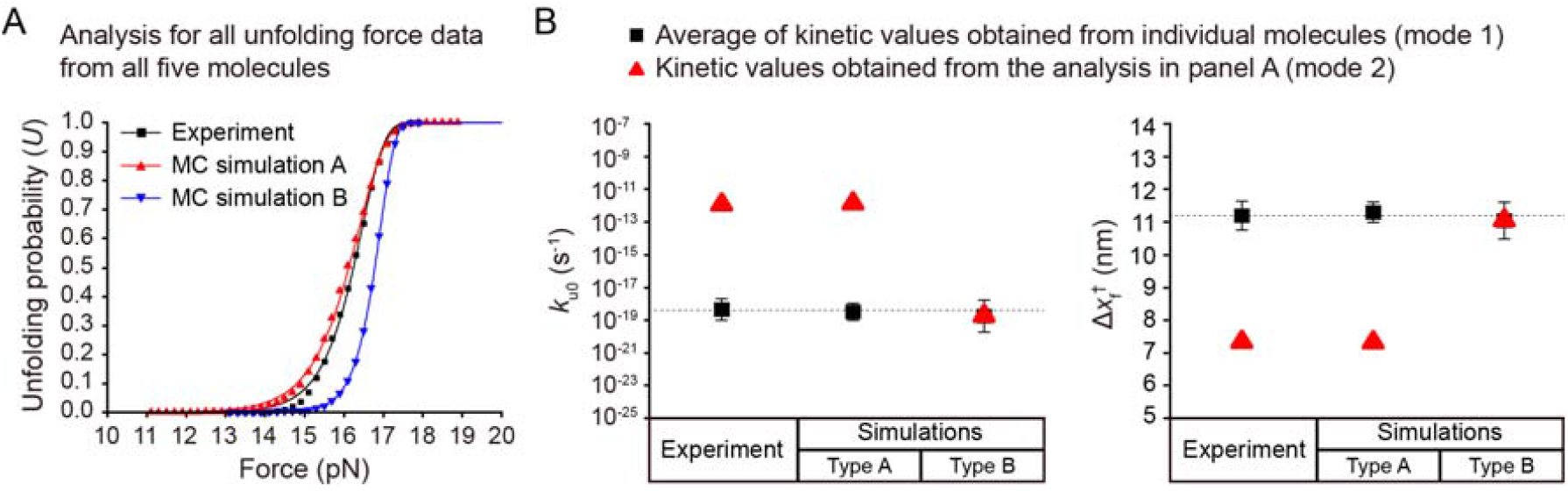
Comparison of two modes of kinetic analyses. (A) Unfolding probabilities as a function of force for the DNA hairpin that were obtained from all unfolding force data from all molecules (*N* = 5 molecules, *n* = 7500 data points; 1500 data points for each molecule). The plots from the experiment and MC simulation-type A (with bead size error) are deviated from that of the MC simulation-type B (without bead size error). The deviations caused by the error of bead size yield wrong kinetic values, as shown in panel B. (B) Kinetic values obtained from the two modes of kinetic analyses. The analyzed kinetic parameters are the unfolding rate constant at zero force (*k*_u0_) and the distance between the native and transition states (Δx_f_^†^). The black squares indicate the average of kinetic values obtained from individual molecules (mode 1). The red triangles indicate the kinetic values obtained from the analysis for all data together, as shown in panel A (mode 2). In the simulation-type B, the obtained kinetic values are identical regardless of the modes of kinetic analyses. For the analysis mode 1, the results from the experiment and simulation-type A match that of the simulation-type B.

**Figure 5–figure supplement 3.**
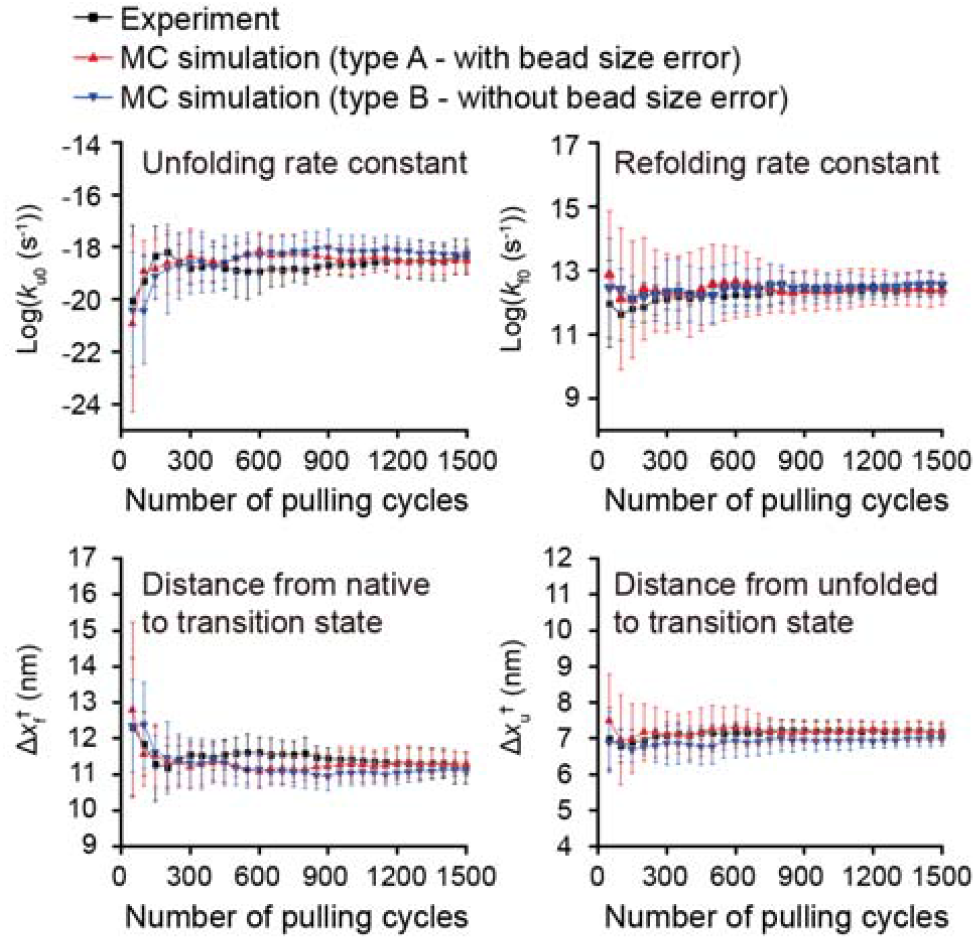
Unfolding and refolding kinetic traces for the DNA hairpin as a function of the number of pulling cycles. The data were obtained from the experiment and MC simulations (*N* = 5 molecules, *n* = 1500 data points). The *k*_u0_ (*k*_f0_) and Δx_f_^†^ (Δx_u_^†^) indicate the kinetic parameters, *i*.*e*., the unfolding (refolding) rate constant at zero force and the distance between the native (unfolded) and transition states, respectively.

**Figure 5–figure supplement 4.**
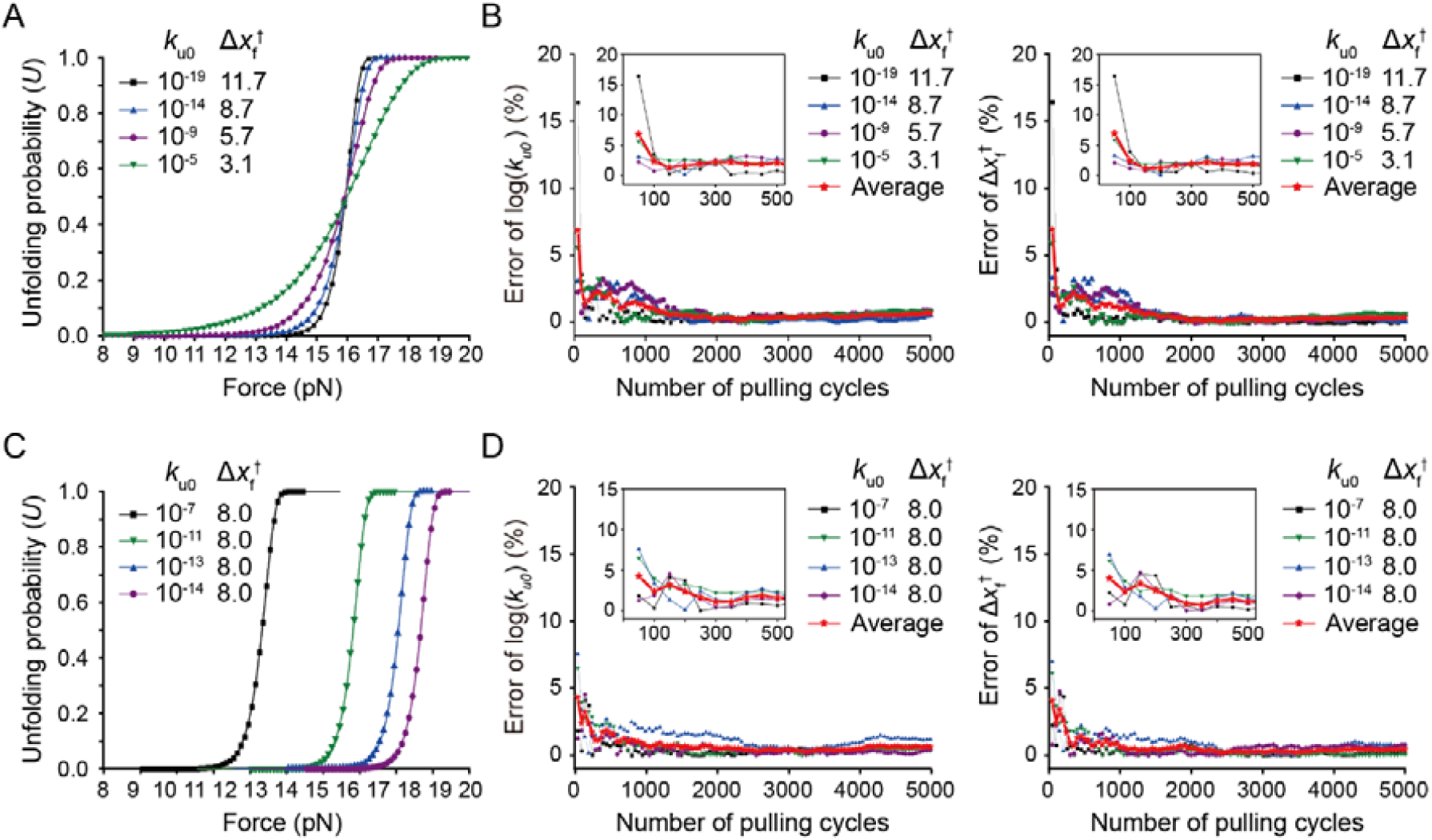
Evaluation of kinetic errors for various shapes of unfolding probability profiles. (A,B) Kinetic errors for unfolding probability profiles with the same *F*_*U*=0.5_ and different slopes. For each set of kinetic values, the panels A and B show the unfolding probability as a function of force (A) and the kinetic error percentage as a function of the number of pulling cycles (B), respectively. (C,D) Kinetic errors for unfolding probability profiles with the same slope and different *F*_*U*=0.5_. For each set of kinetic values, the panels C and D show the unfolding probability as a function of force (C) and the kinetic error percentage as a function of the number of pulling cycles (D), respectively. All the data were created from the MC simulation-type B, the results of which match those of the experiment in the analysis mode 1, as shown in Figure 5–figure supplement 2. In the panels B and D, the reference point of zero error is the one for 10000 times of the pulling. The error trace for each parameter set is the averaged one for the data from all molecules at each cycle (*N* = 10). The red curves are the averaged one for the averaged kinetic traces at each cycle, which are shown as individual traces in Figure 5.

## Source Data and Codes

Figure 1–source data 1. Original files of raw unedited gels and figures with uncropped gels.

Figure 3–source data 1. Full list of tested conditions for DBCO-azide conjugation.

Figure 4–source data 1. Survey of protein unfolding forces in optical and magnetic tweezers.

Figure 4–source data 2. Raw data for the figure panels.

Figure 5–source data 1. Raw data for the figure panels.

Figure 5–source data 2. Codes for the Monte Carlo simulations

**Figure 3–source data 1.**
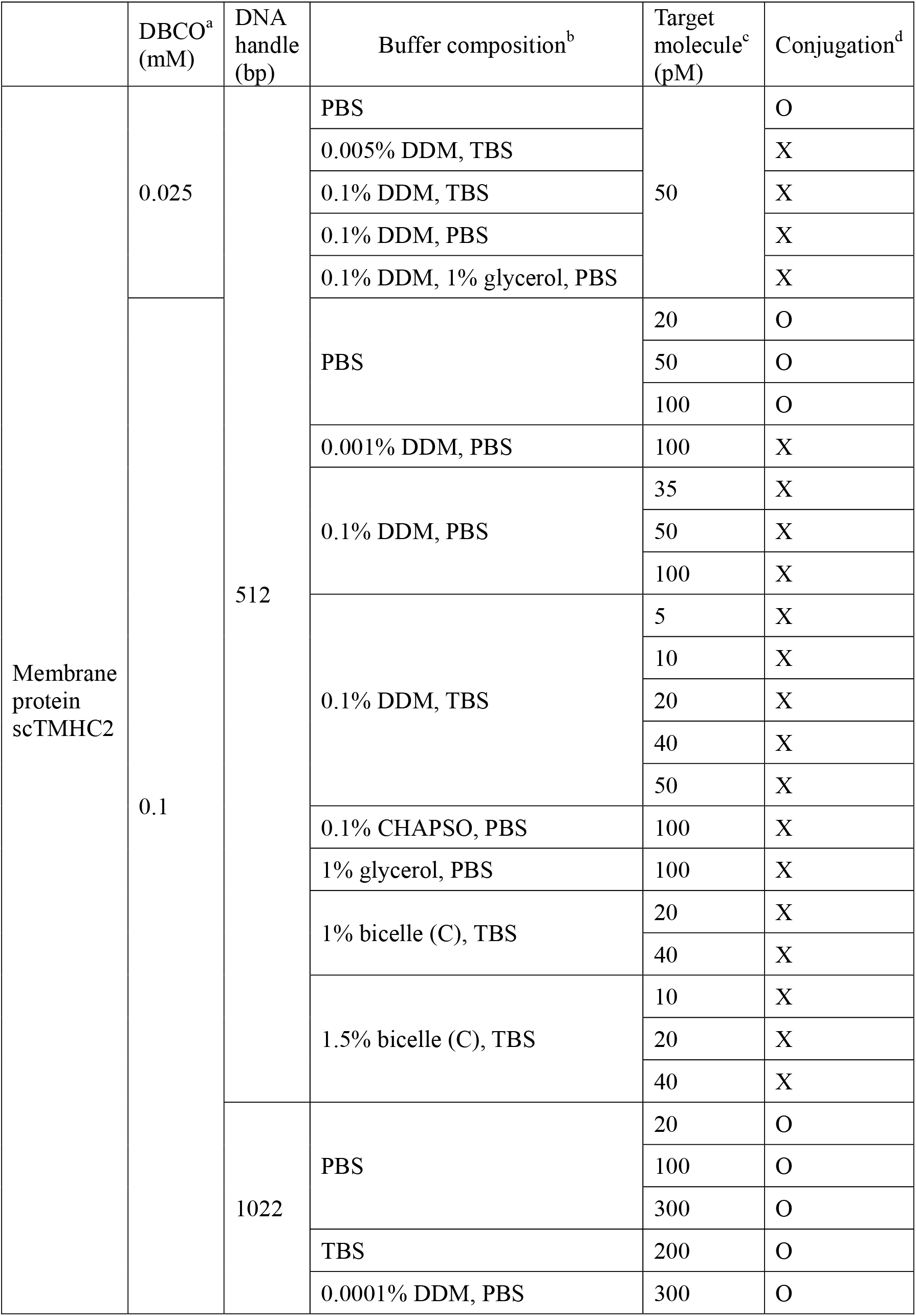
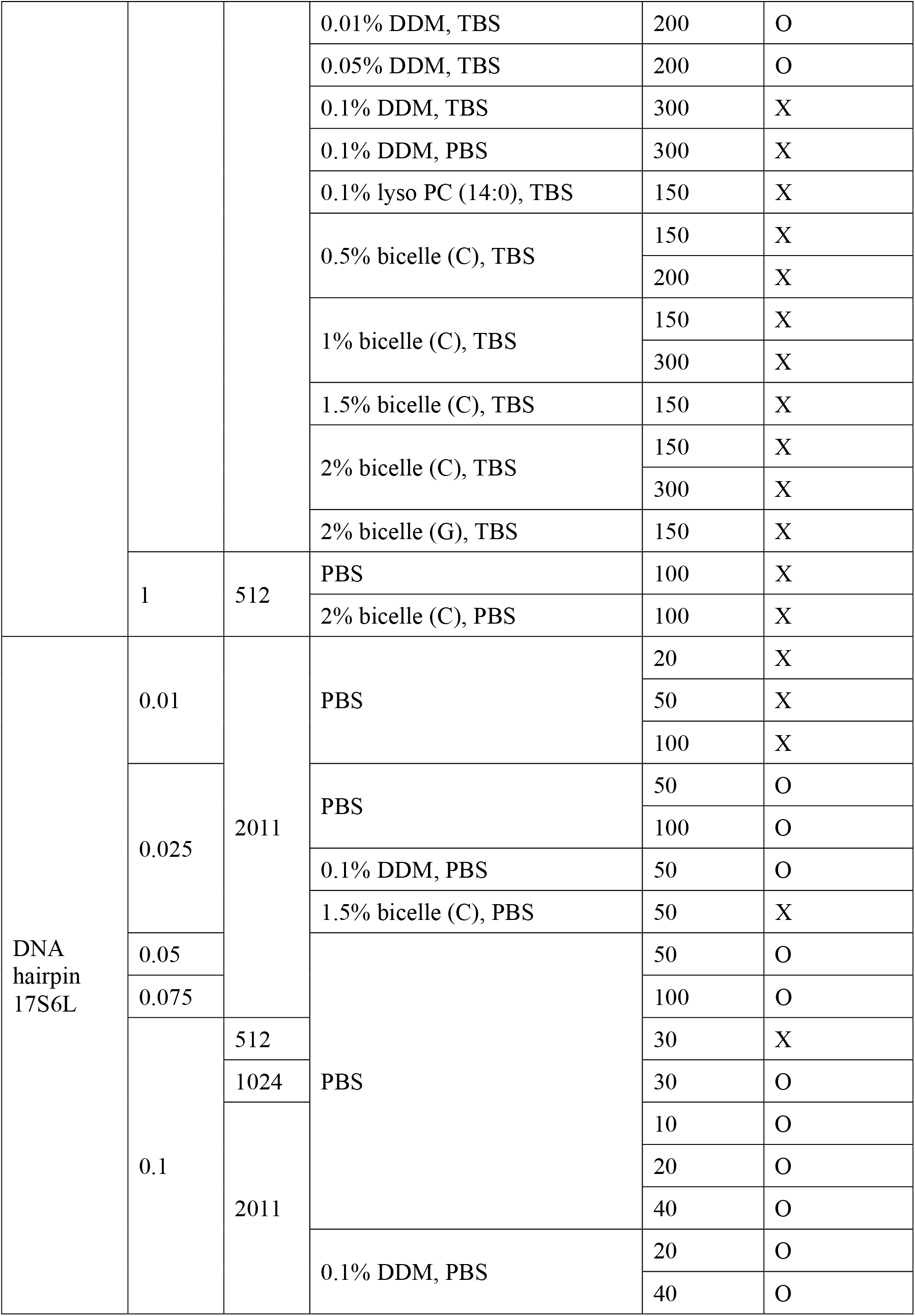
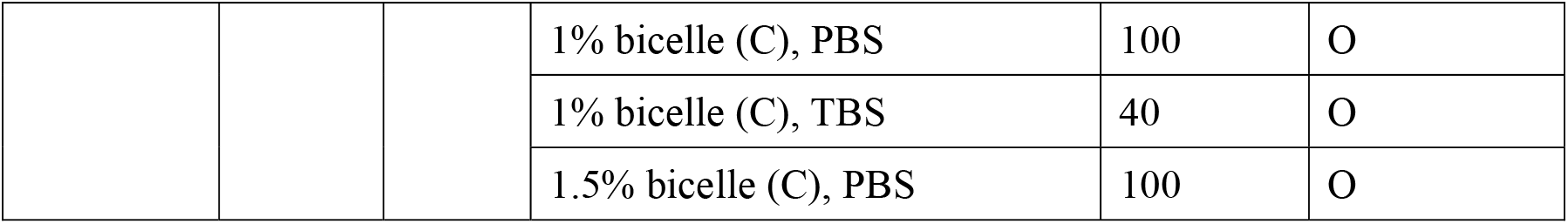
Full list of tested conditions for DBCO-azide conjugation. ^a^ DBCO concentration indicates the final concentration of DBCO-sulfo-NHS crosslinker in the reaction of DBCO modification on bead surface. Others (buffer composition, DNA handle, targe molecules) indicate the conditions in the DBCO-azide conjugation reaction in single-molecule sample chambers. ^b^ PBS indicates a phosphate-buffered saline (0.1 M sodium phosphate, 150 mM NaCl, pH 7.3). TBS indicates a tris-buffered saline (50 mM Tris-HCl, 150 mM NaCl, pH 7.5). Bicelle (C or G) is composed of DMPC (or DMPG) lipid and CHAPSO detergent at a 2.5:1 molar ratio. The % indicates w/v %. ^c^ Final concentration of target molecules (the membrane protein or DNA hairpin) in the tethering to the sample chamber surface. ^d^ The symbol O indicates successful conjugation between DBCO-modified magnetic beads and target membrane proteins so that specifically-tethered beads can be found and force-extension curve data can be obtained (confirmed by three replicates). The symbol X indicates unsuccessful conjugation to target proteins or entire nonspecific binding of beads to the chamber surface. In this case, no data is obtained.

Refer to the source data file, Figure 4-source data 1, for the references in this table

**Figure 4-source data 1.**
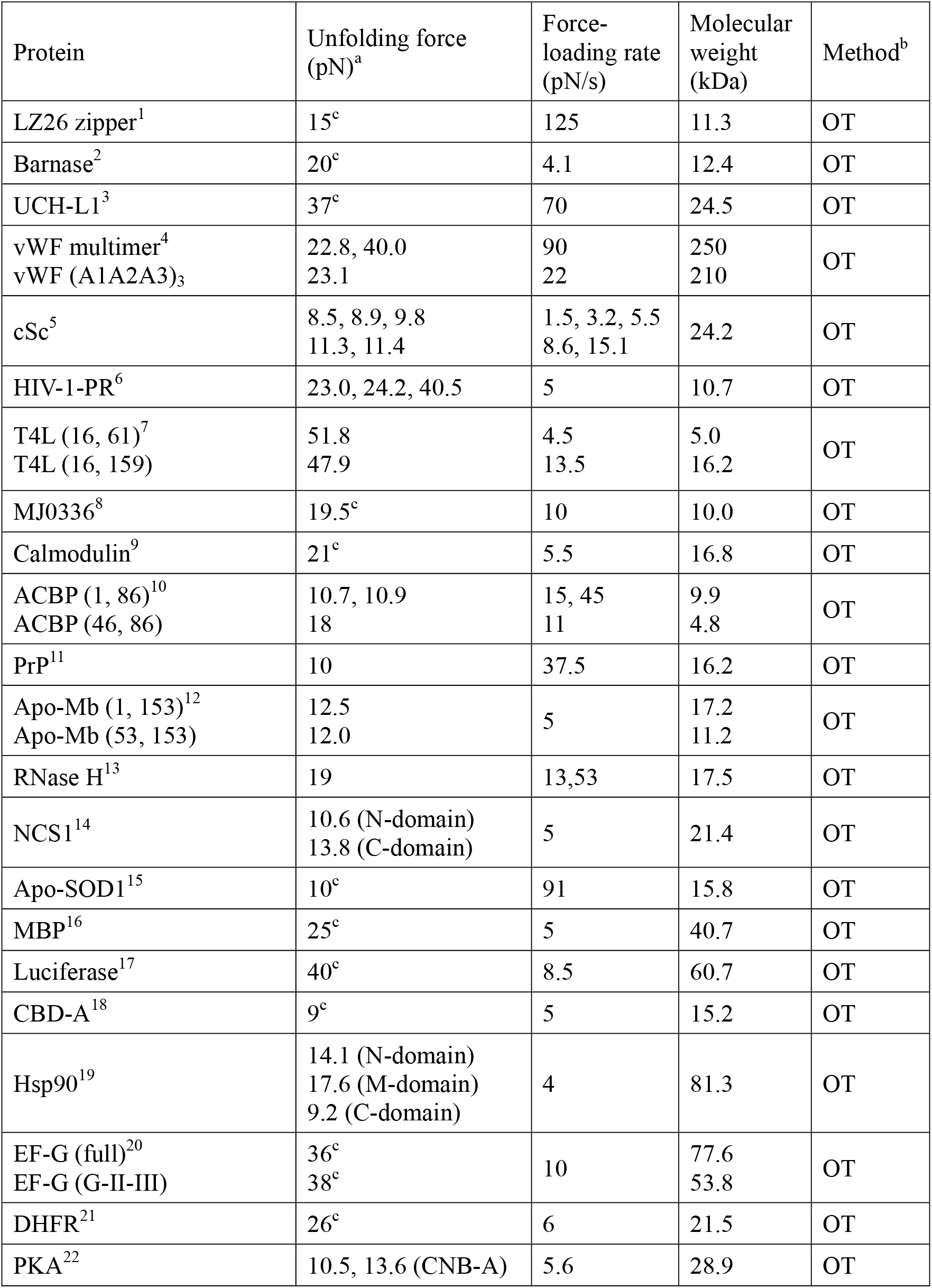
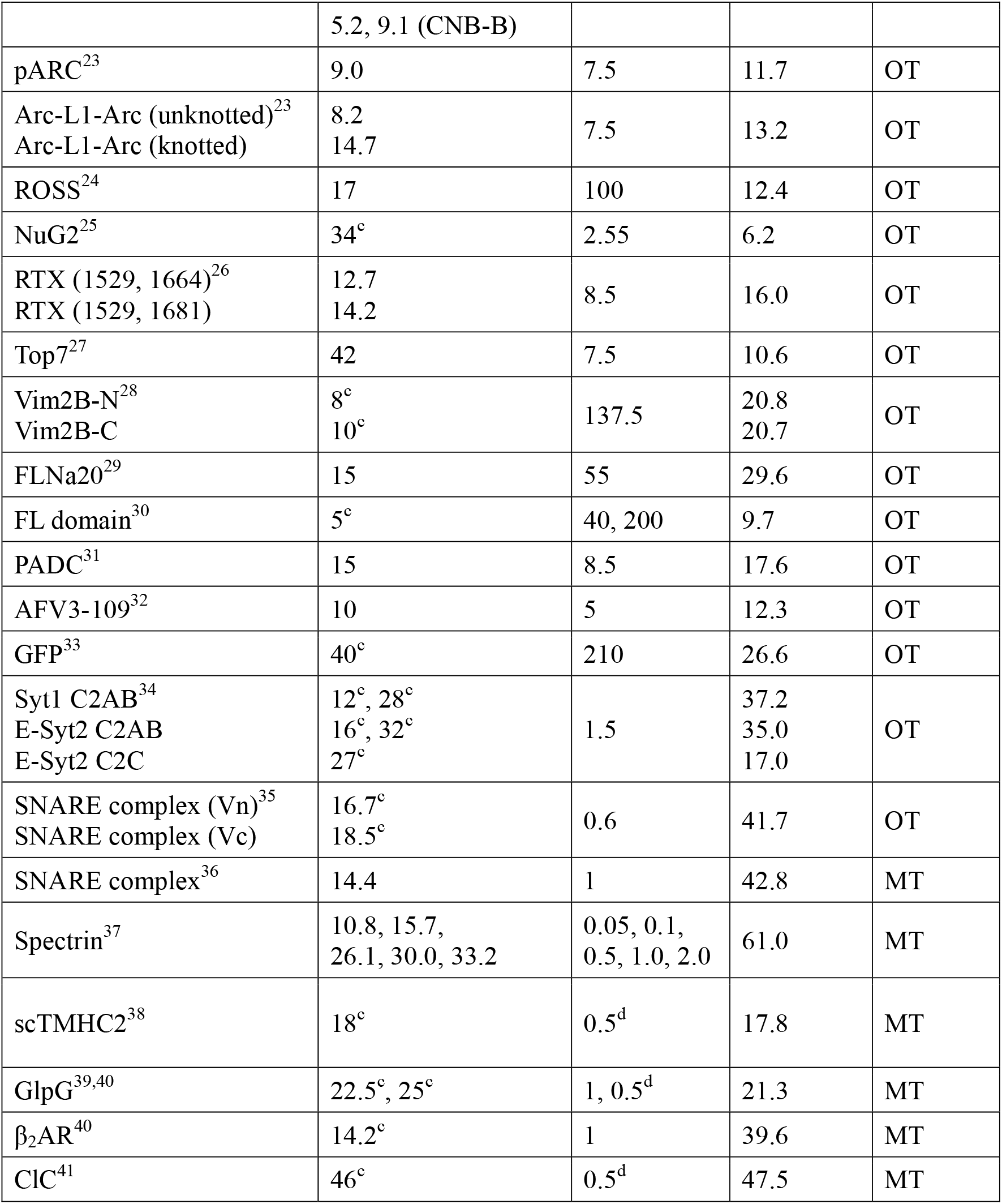
Survey of protein unfolding forces in optical and magnetic tweezers. ^a^ The unfolding force indicates the most probable unfolding force in unfolding force distribution except the items with the superscript c. ^b^ OT and MT indicate optical tweezers and magnetic tweezers, respectively. ^c^ The unfolding force is an approximate or averaged value. ^d^ The force-loading rate is an averaged value over the force scanning of 1-50 pN.

